# Gene expression profiling reveals subgenome dominance during *Brassica napus* seed development

**DOI:** 10.1101/2020.04.29.068189

**Authors:** Deirdre Khan, Dylan J. Ziegler, Jenna L. Kalichuk, Vanessa Hoi, Nina Hyunh, Abolfazl Hajihasani, Isobel A.P. Parkin, Stephen J. Robinson, Mark F. Belmonte

**Author notes:** Author Contributions: DK, JLM, NH, VH, and AH performed experiments. IAPP and SR provided the DH12075 genome and annotation. DK, DJZ, SJR, IAPP and MFB prepared the manuscript. All authors read and approved the final manuscript. MFB agrees to serve as the author responsible for contact and ensures communication.

## Abstract

We profiled the gene regulatory landscape of *Brassica napus* reproductive development using RNA sequencing. Comparative analysis of this nascent amphidiploid across the plant lifecycle revealed the contribution of each subgenome to plant reproduction. Global mRNA profiling revealed lower accumulation of C^n^ subgenome transcripts relative to the A^n^ subgenome. Subgenome-specific transcriptional networks identified distinct transcription factor families enriched in each of the A^n^ and C^n^ subgenome early in seed development. Global gene expression profiling of laser-microdissected seed subregions further reveal subgenome expression dynamics in the embryo, endosperm, and seed coat of early stage seeds. Transcription factors predicted to be regulators encoded by the A^n^ subgenome are expressed primarily in the seed coat whereas regulators encoded by the C^n^ subgenome were expressed primarily in the embryo. Data suggest subgenome bias are characteristic features of the *B. napus* seed throughout development, and that such bias might not be universal across the embryo, endosperm, and seed coat of the developing seed. Whole genome transcription factor networks identified BZIP11 as a transcriptional regulator of early *B. napus* seed development. Knockdown of *BZIP11* using RNA interference resulted in a similar reduction in gene activity of predicted gene targets, and a reproductive-lethal phenotype. Taken together, transcriptional networks spanning both the A^n^ and C^n^ genomes of the *B. napus* seed can identify valuable targets for seed development research and that-omics level approaches to studying gene regulation in *B. napus* can benefit from both broad and high-resolution analyses.

**One Sentence Summary:** Global RNA sequencing coupled with laser microdissection provides a critical resource to study subgenome bias in whole seeds and specific tissues of polyploid plants.

## Introduction

*Brassica napus* (canola) is the second most important oilseed crop in the world with global production of 75 million tons in 2017 (FAOSTAT 2017). It is a relatively recent allopolyploid derived from hybridization between *B. rapa* and *B. oleracea* (Nagaharu, 1935). The resulting plant is an amphidiploid species which has retained its progenitor subgenomes from *B. rapa* (A^r^A^r^) and *B. oleracea* (C°C°) to form *B. napus* (A^n^A^n^C^n^C^n^). Despite its importance to the global economy, *B. napus* genome dynamics are poorly understood, especially in the context of gene regulation during seed development. The developmental transition from seed morphogenesis to maturation (Figure 1A) requires a re-orchestration of the genes and gene regulatory networks underpinning seed development. Therefore, *B. napus* represents an appropriate model for research on transcriptomic changes across seed development of economically important polyploid plants.

**Figure 1.**
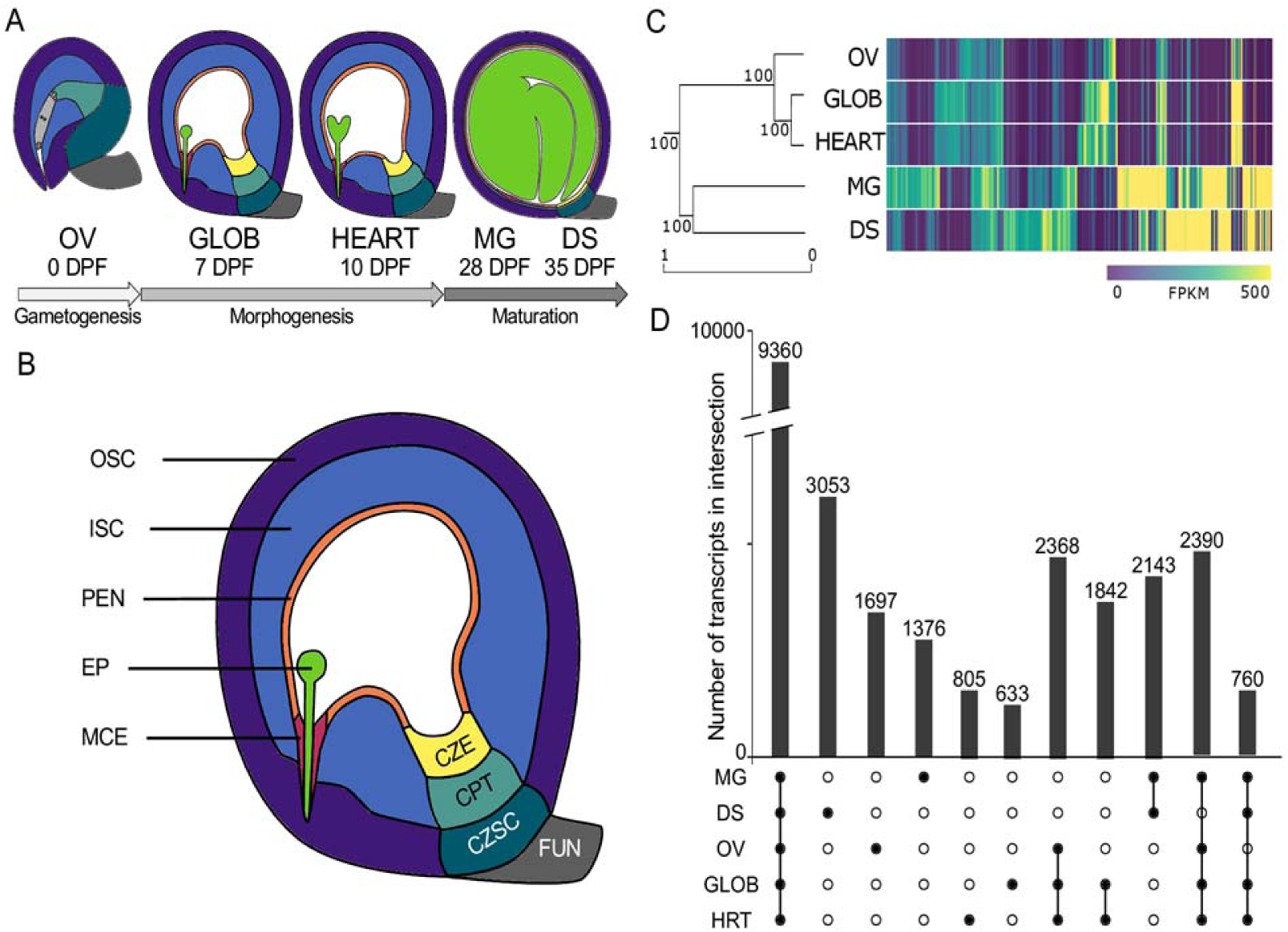
The dynamic transcriptome of *Brassica napus* seed development. (A) Stages of seed development examined in study. *Brassica* napus cv DH12075 tissue schematic. (B) The globular stage seed is expanded, showing the outer distal seed coat (OSC, indigo), inner distal seed coat (ISC, blue) chalazal seed coat (CZSC, dark teal), chalazal proliferating tissue (CPT, light teal), chalazal endosperm (CZE, yellow), peripheral endosperm (PEN, orange), micropylar endosperm (MCE, pink), embryo (EP, green), and funiculus (FUN, grey). (C) hierarchical clustering analysis of seed developmental stages: ovule (OV), globular (GLOB), HEART, mature green (MG), and dry seed (DS) paired with a heatmap of the top 1000 most highly expressed genes. High level of expression (FPKM>500) is indicated in yellow. (D) Upset plot showing intersections of transcript accumulation among the OV, GLOB, HRT, MG, and DS seed stages. Intersections of black circles are analogous to a union on a Venn diagram. Black circles with no intersections indicate a population of exclusive transcripts. All data used for these analyses are found in Dataset S1.

*Brassica napus* is a nascent amphidiploid which has not yet undergone substantial fractionation, and consequently has inherited two distinct diploid subgenomes one from each of its progenitors (Chalhoub et al., 2014). However, the A^n^ and C^n^ subgenomes of *B. napus* are still divergent from their progenitor species, with numerous chromosomal translocations and inversions (Zou et al., 2016; Hurgobin et al., 2018) and novel transposable elements (Yang et al., 2016) present between the progenitor species and their amphidiploid hybrids. This neo-polyploidy presents an opportunity to examine gene regulation in polyploid plants in their infancy. The current question of how two distinct subgenomes are regulated to orchestrate seed development in *B. napus* is further complicated by the potential interdependence of the two subgenomes. Many polyploid species display evidence of subgenome expression dominance and/or fractionation (Cheng et al., 2018). Gene loss in the subgenomes of *B. napus* has occurred more quickly than in the progenitor *B. rapa* and *B. oleracea* (Chalhoub et al., 2014), which may also allow for specialization of subgenomes. Moreover, homoeologous exchanges have occurred between the two subgenomes (Hurgobin et al., 2018), further complicating interactions or interdependencies between the two subgenomes.

Expression bias favouring the *B. rapa* subgenome has been observed in the leaves of newly resynthesized *B. napus* (Wu et al., 2018) indicating that expression bias is observable immediately after allopolyploidization and continues in subsequent generations. Comparisons of newly synthesized and extant allopolyploids of monkeyflower (*Erythranthe peregrina* G.L. Nesom) also indicate that expression bias occurs immediately after polyploidization but also that it increases across subsequent generations. Studies in *Senecio* (Hegarty et al., 2006) and *Spartina* (Chelaifa et al., 2010) also record large scale transcriptional shifts following hybridization. Thus, we must consider both subgenome-enriched and whole genome contributions to gene regulatory networks in *B. napus* seed development. *Brassica napus* shows expression bias in its vegetative tissues as well as tissue-specific effects on expression bias between leaves and roots (Chalhoub et al., 2014). Since there are observable expression biases among vegetative and reproductive tissues in *B. napus*, we seek to also understand if there is observable subgenome expression bias in specific tissues of the developing seed.

The seed itself is composed of three ontogenetically distinct tissues: the maternal seed coat, the zygotic endosperm and the embryo. All three tissues undergo dramatic developmental changes across the seed lifecycle (Figure 1A). Even in the model plant Arabidopsis, our understanding into the communication between these tissues is still lacking and only recently has begun to be discovered (Nowack et al., 2010; Belmonte et al., 2013; Figueiredo and Köhler, 2014; Xu et al., 2016). Such dramatic changes in transcript profiles are particularly complex in neo-allopolyploids like *B. napus* which must also regulate two distinct subgenomes during complex developmental processes. Recently, we examined the tissue-specific transcriptome landscape of the globular stage *B. napus* seed (Ziegler et al., 2019), and now have the ability to examine the contributions of each subgenome across seed development in both filial and maternal tissues.

In the current paper, we present a global transcript analysis and functional characterization of genes required to program the *B. napus* seed. Transcription factor circuits defining seed development and maturation further deepen our characterization of the seed. We observe subgenome bias that favours the expression of the A^n^ subgenome across all stages of seed development, as well as among subregions of the early stage seed. We also identify transcriptional regulators that define the transcript accumulation patterns of both the A^n^ and C^n^ subgenomes. Whole genome transcription network analyses presented in this work predict transcription factor activity essential to seed development. Functional characterization of these networks provides further evidence into the complexity of TF-promoter interactions that program the making of a seed.

## Results

### Seed development is characterized by global shifts in gene activity

We profiled gene activity throughout the genome across five stages of reproductive seed development in *B. napus* (Figure 1A). Clustering analysis using all detected transcripts showed that each stage of seed development is transcriptomically distinct. Furthermore, early (OV, GLOB, HEART) and late (MG, DS) seed stages form distinct groups (Figure 1C). In total, we detected 32,712 transcripts across seed development, 9,360 of which were expressed in all stages studied (FPKM > 5, Figure 1D, Figure S1, Dataset S1).

Differential gene expression analysis revealed stage- and phase-enriched transcripts that accumulate in the developing *B. napus* seed (FPKM>10 in stage/phase, FPKM<10 in other stages/phases) and recapitulate our clustering analysis (Figure 1D, Dataset S2). There is strong transcriptional continuity between pre- and post-fertilization seed development: the early stages of reproductive development (OV, GLOB, HEART) share more transcripts (2,368) than the morphogenesis phase alone (GLOB and HEART, 1,842 transcripts). The OV stage (1,697) harbours more than double the number of unique transcripts than either the GLOB (805) or HEART (633) stages. The dry seed is most unique and contains the greatest number of specific transcripts compared to other seed stages (Figure 1D, Dataset S2).

Next, we were interested in studying differentially accumulating transcripts in each of the A^n^ and C^n^ subgenomes in the seed. 38.2% of all annotated A subgenome-encoded transcripts were expressed, compared to 30.7% of transcripts encoded by the C subgenome (Dataset S1, FPKM>5). Considering all data points, transcripts encoded by the A^n^ subgenome accumulated to a greater extent than those encoded by the C^n^ subgenome (Figure 2A, p<0.001, Mann-Whitney-Wilcoxon). Transcript accumulation bias towards the A^n^ subgenome is significant across the seed lifecycle and was more apparent in the early phase of seed development (OV, GLOB, HEART).

**Figure 2.**
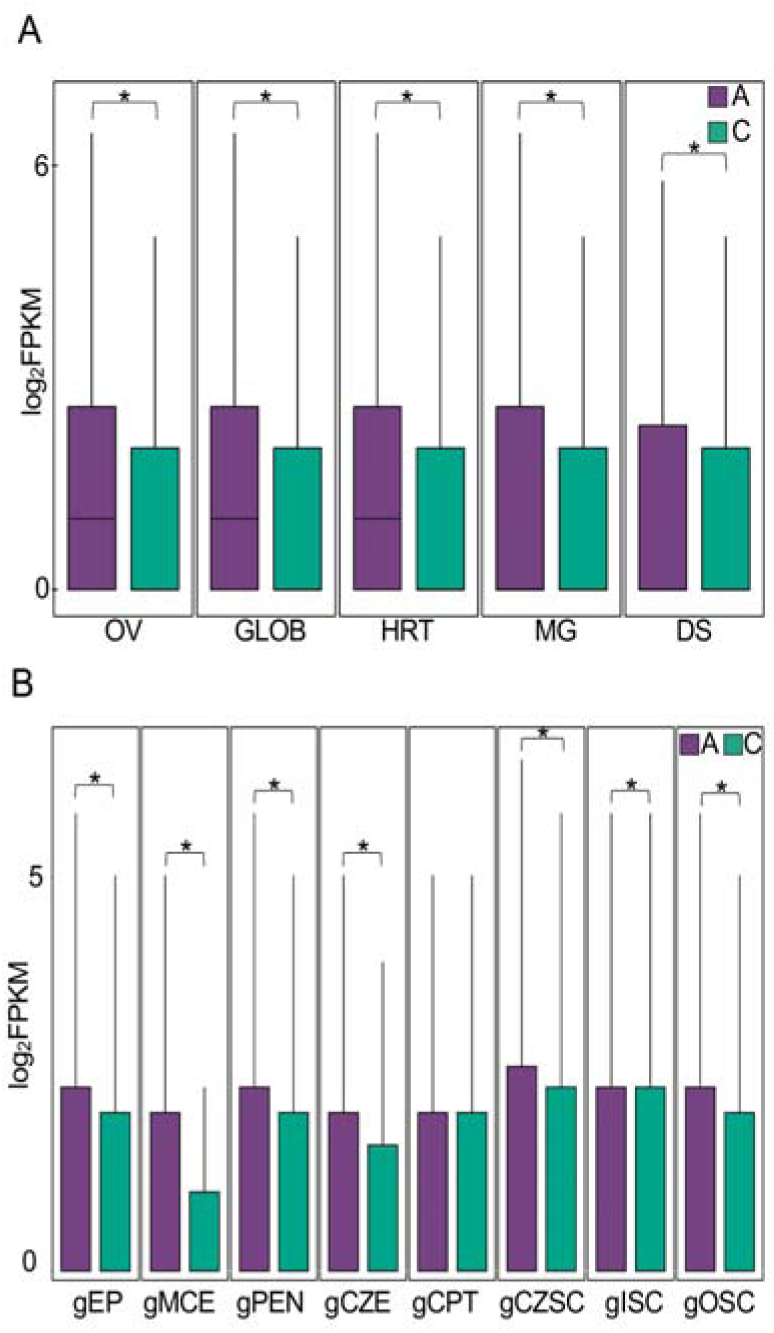
Transcript accumulation from the A^n^ and C^n^ subgenomes in developing *Brassica napus* seeds. (A) Quantile boxplots of all detected transcripts from the A^n^ (purple) and C (green^n^ () subgenomes in all seed stages. (B) Quantile boxplots of all detected transcripts from the A^n^ and C^n^ subgenomes in each seed tissue at the globular stage of seed development. EP = Embryo Proper, MCE = Micropylar Endosperm, PEN = Peripheral Endosperm, CZE = Chalazal Endosperm, CPT = Chalazal Proliferating Tissue, CZSC = Chalazal Seed Coat, ISC = Inner Seed Coat, OSC = Outer Seed Coat. Data used to identify expression dominance between the two subgenomes are found in Dataset S1.

We then examined micro-dissected region-specific seed expression data from the GLOB stage of *B. napus* seed development, schematic in Figure 1B). Of all known transcripts, 48% of A subgenome coded transcripts, and 42% of C subgenome encoded transcripts accumulated across the GLOB seed regions. At the GLOB stage of seed development, higher transcript accumulation from the A^n^ subgenome is significant in every region except for the chalazal proliferating tissue (CPT) (Figure 2B, p<0.001, Mann-Whitney-Wilcoxon). Therefore, at the GLOB stage of development, the embryo, endosperm, and seed coat regions (except the CPT) all contribute to higher accumulating transcripts of the A^n^ subgenome.

### Seed maturation is characterized by the high expression of lipid and protein storage genes

The most highly expressed transcripts in our dataset accumulate in the maturation phase of seed development (Figure 1C). In particular, the MG stage accumulated high transcript levels of seed storage albumin (*BnSESA1, BnSESA3, BnSESA4*), oleosin (*BnOLEO1, BnOLEO2, BnOLEO3*), and cruciferin (*BnCRU1, BnCRU2, BnCRU3*) encoding genes. The DS stage was characterized by the high expression of LATE EMBRYOGENESIS ABUNDANT PROTEIN (*BnLEA, BnLEA14, BnLEA18, BnLEA4-5*) and dehydrin (*BnRAB18*) encoding genes (Figure S2). Network analysis of MG-enriched transcripts derived from the A^n^ and C^n^ subgenomes yielded dramatically different predictions of regulatory circuits in seed maturation (Figure S3). While both networks predict regulators of seed maturation, glucosinolate metabolism, and organ positioning, the A^n^ subgenome network predicts numerous MYB and bHLH transcription factors as regulators while the C^n^ subgenome network predicts only FUS3 and WOX12 (via the homeodomain binding motif) as transcriptional regulators (Figure S3). Our analysis found a more intact maturation network in the A^n^ subgenome than observed in the C^n^ subgenome. There is no transcriptome data yet available for the MG stage or other maturation stages of *B. napus* seed that compares the different subregions of the seed coat, endosperm and embryo. Thus, it is not clear how each region contributes to the observed differences in each of the subgenomes during the maturation phase of seed development.

### Transcription factor networks predict distinct contributions of the A^n^ and C^n^ subgenomes to the regulation of early *B. napus* seed development

We were curious to see if there were temporally distinct regulatory networks operating within the A^n^ and C^n^ subgenomes of the developing *B. napus* seed. We split the temporally distinct genes by subgenome of origin and performed network enrichment analysis to predict transcription factor binding and gene ontology enrichment. This analysis predicts networks of transcription factors, DNA binding motifs, and expressed genes only within a single subgenome. Isolating the subgenomes in this way assumes that there is no cross-talk between them, which is unlikely to represent the reality of gene regulatory networks in the seed. However, this does allow us to determine if intact regulatory networks can still be detected if we separate the transcriptome by subgenome of origin.

Transcriptional subgenome bias is more evident in the early stages of seed development (Figure 2A). Transcription factor networks derived from both the A^n^ and C^n^ subgenomes during early seed development predicted transcriptional regulation of several common biological processes including cell division/cell cycle, ovule development, and histone phosphorylation (Figure 3A-B). Our analysis predicted homologues from both A^n^ and C^n^ subgenomes of MYB17, MYB32, MYB61, and MYB95 as regulators of early seed development. The A^n^ subgenome transcriptome is enriched for MADS box motif-binding transcription factors canonically associated with ovule, flower and maternal seed development such *as TRANSPARENT TESTA16* (TT16) and *SEPELLATTA1* (SEP1), *SEEDSTICK* (*STK*), and *SEPELLATA2* (*SEP2*) (Figure 3A). The early seed C^n^ subgenome transcription factor networks were enriched for bHLH and BES TFs that were absent in the A^n^ subgenome network (Figure 3B).

**Figure 3.**
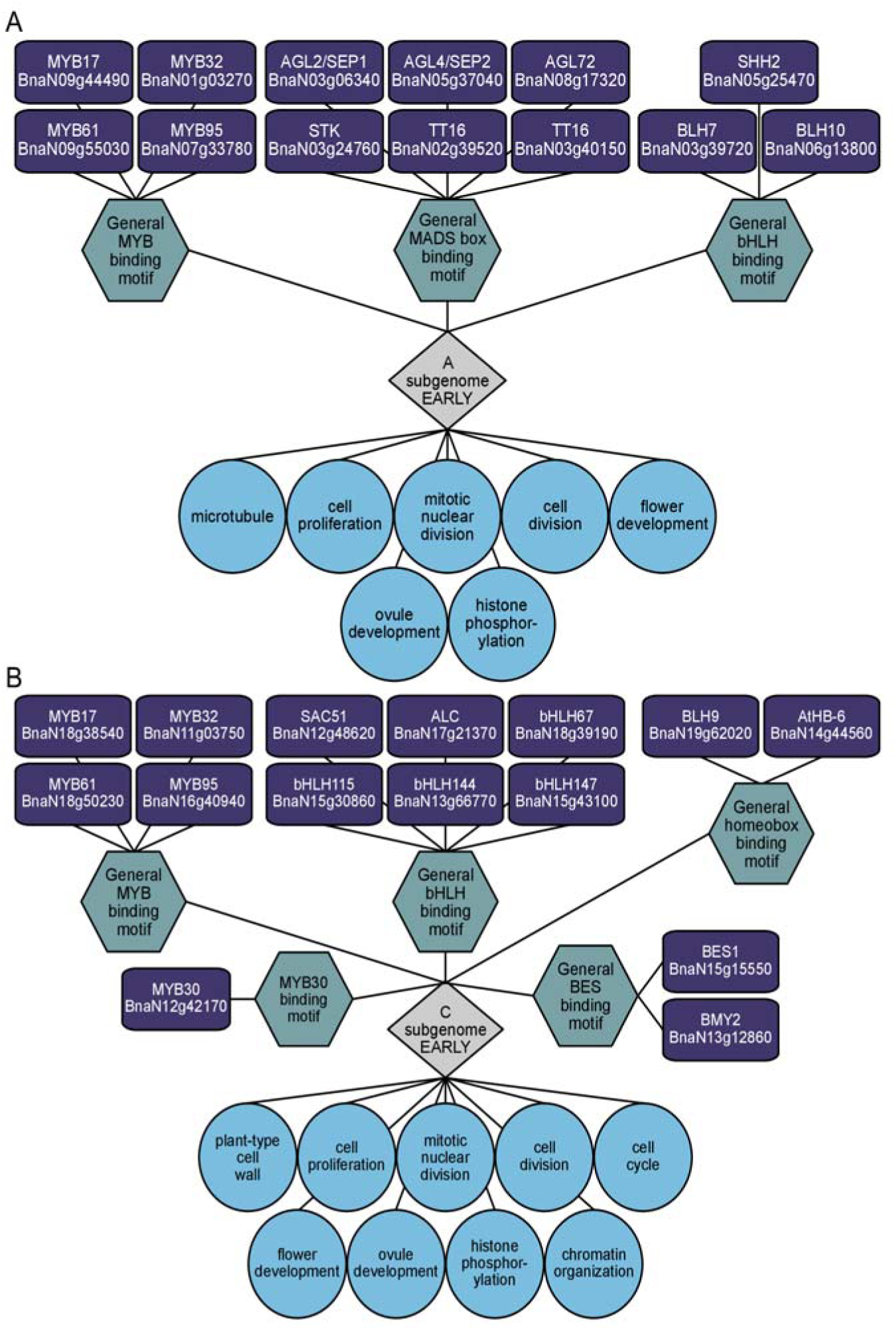
Predictive regulatory networks of early *B. napus* seed development. Genes belonging to a co-expression network (grey diamond) are enriched for transcription factors (purple rectangles) predicted to bind to DNA sequence motifs (teal hexagons) upstream of genes associated with biological processes (blue circles). (A) Network of early phase coexpressed genes originating in the A^n^ subgenome. (B) Network of early phase coexpressed genes originating in the C^n^ subgenome. For a detailed explanation co-expression networks generated by SeqEnrich, see Becker et al., 2017. Clustering outputs, coexpressed genes, and enrichment outputs can be found in Dataset S2.

We then examined available tissue-specific transcriptome data for the GLOB stage *B. napus* seed (Ziegler et al., 2019) to pinpoint how each subregion contributed to the A^n^ and C^n^ subgenome networks (Figure 4). Spatial transcriptomic analysis of the early *B. napus* seed confirms a strong maternal contribution from the A^n^ subgenome to the MADS box, MYB and homeobox clades. Only three of the A^n^ subgenome enriched transcription factors show expression in filial subregions and are expressed in the micropylar endosperm. The filial subregions, in particular the embryo proper and micropylar endosperm, show greater contribution of the C^n^ subgenome enriched TFs, especially in the bHLH and BES binding clades Thus, we can detect maternal transcriptional circuits in the A^n^ subgenome of the early *B. napus* seed, and find distinct C^n^ subgenome contributions to the filial transcriptome.

**Figure 4.**
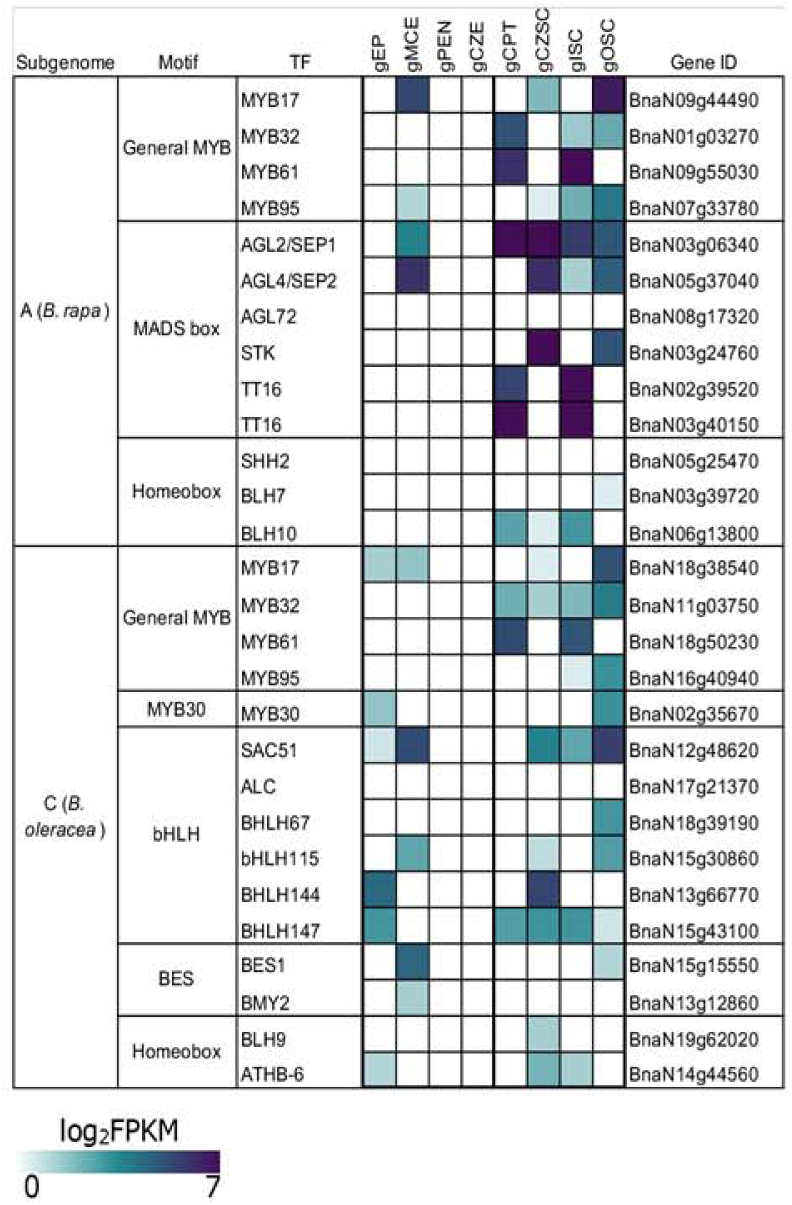
Heatmap of transcription factor expression levels in the different tissues of the GLOB stage seed. (A) Tissue schematic of a globular stage seed. (B) Tissue-specific gene activity of transcription factors identified in Figure 3. FPKM values have been log2 transformed. Darker blue colour represents increased transcript abundance.

### BZIP transcription factors play a role in the regulation of morphogenesis phase seed development in *B. napus*

We first examined the A^n^ and C^n^ transcriptomes together in order to identify transcription factors that may contribute to gene regulation in seed development. Enrichment analysis of co-expressed gene sets (Fuzzy K means clustering) revealed transcription factor networks in the *B. napus* seed (Figure 5A). Co-expressed genes up-regulated early (OV, GLOB, HRT) identified three Group S1 BZIPs (*BnBZIP11, BnBZIP44, BnBZIP53*) and one Group C BZIP (*BnBZIP25*) as regulators of photosynthetic and cytoskeletal processes via the binding of a general BZIP-binding DNA sequence motif (Figure 5A). Previously sucrose-mediated translational repression was found to be enriched in morphogenesis phase transcripts, a phenomenon known to modulate translation of *BZIP11* and *BZIP44* (Rahmani et al., 2009). We chose to investigate *BnBZIP11* as a target gene for further study given its potential role in modulating sugar (Rahmani et al., 2009; Ma et al., 2011), auxin (Weiste et al., 2014; Weiste et al., 2017) and amino acid signalling (Hanson et al., 2008) in the chalazal pole of the early *B. napus* seed (Ziegler et al., 2019). We generated knockdown mutants of *BnBZIP11* (*BnaN08g13560, BnaN11g74680, BnaN13g74680*) using RNA interference in *Brassica napus* (cv. Topas). Transcript levels of *BnBZIP11* were reduced 99.5% in the T_1_ heart stage mutant siliques (Figure 5B). These mutants exhibited a reduced seed set with 53% reproductive lethality (Figure 5C, Student’s T-Test, p<0.05). To test if reduced abundance of *BZIP11* leads to reduced BZIP11 activity and therefore reduced expression of its target genes, we identified several target genes predicted to be regulated by *BnBZIP11* in the co-expression network (Figure 5A) and quantified transcript levels in the T_1_ mutant siliques (Figure 5B). *ASN1* (*GLUTAMINE-DEPENDENT ASPARAGINE SYNTHASE*) is a known target of BZIP11 (Hanson et al 2007) and was used as a positive indicator. The two detectable homologues of *ASN1* were knocked down >85% in the *BnBZIP11* knockdown mutant (*BnaN13g62650*, 90.9%; *BnaN13g62660*, 85.8%). Two homologues of *BnTUA6* (*TUBULIN ALPHA CHAIN 6*) were also reduced over 80% (*BnaN05g17860*, 85.3%; *BnaN13g09030*, 83.3%), and four homologues of TUB7 were also reduced with variable efficacy (*BnaN04g19540*, 65.4%; *BnaN05g14010*, 55.5%; *BnaN12g31120*, 85.1%; *BnaN14g50530* 62.5%) (Figure 5B).These results suggest that BZIP11 might be at least partially responsible for regulating photosynthesis and cytoskeleton-related gene activity in the early *B. napus* seed.

**Figure 5.**
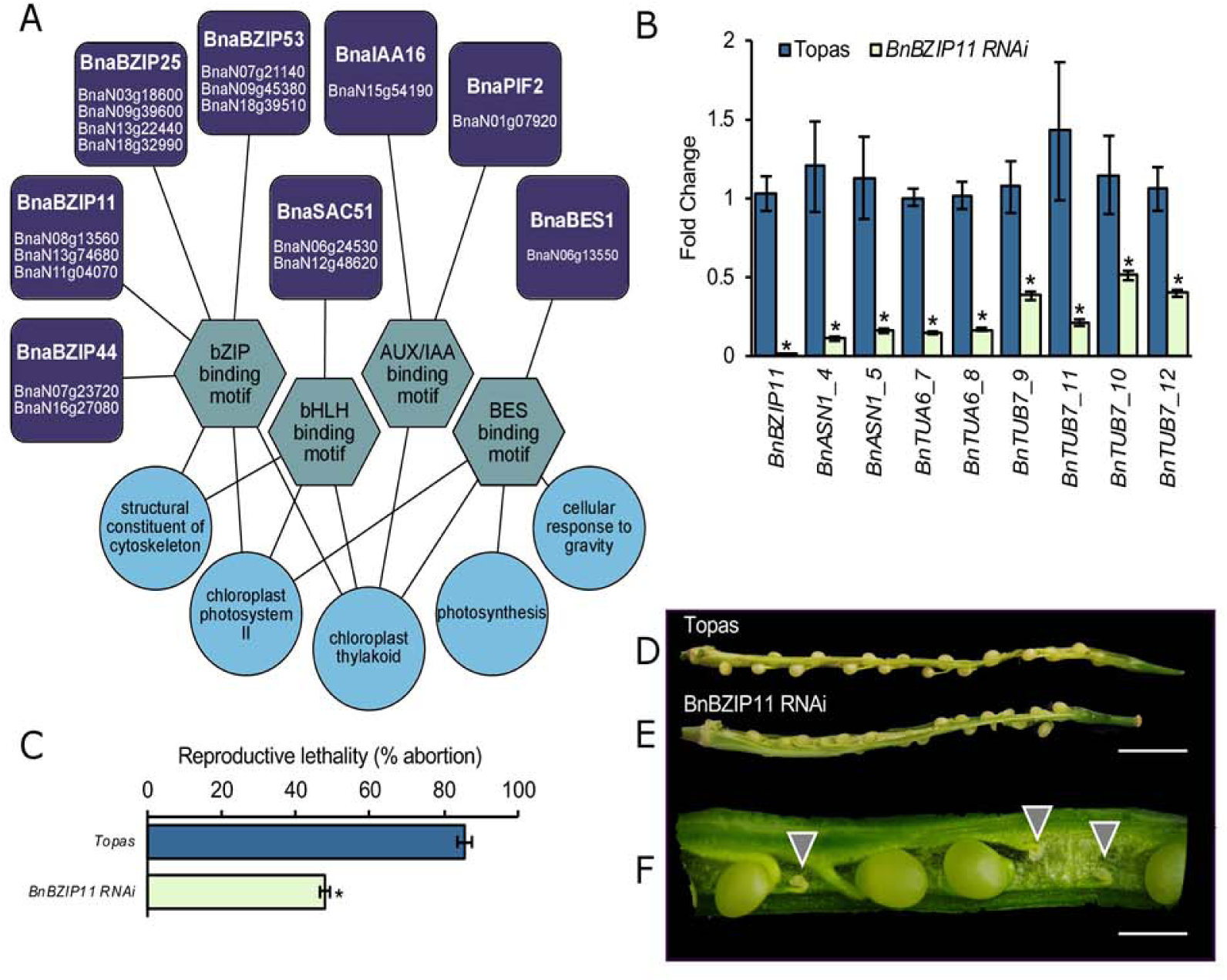
Gene regulation in morphogenesis phase seed development. (A) Putative regulatory network of *B. napus* seed morphogenesis. The network shows transcription factors (purple squares) binding to DNA sequence motifs (teal hexagons) upstream of genes involved in biological processes (GO terms, green circles), all within a set of co-expressed genes during seed morphogenesis. A complete list of the genes associated with this co-expression network as well as all enrichment outputs can be found in Dataset S2 (B) Relative transcript accumulation of *BZIP11* and selected target genes as quantified using qPCR. Targets identified in the GO terms of (A) were selected for expression analysis in *BnBZIP11*_*RNAi*_ mutants. *ASN1*, a known target of BZIP11 was examined. Transcript levels of targets in *BnBZIP11*_*RNAi*_ plants are shown relative to WT Topas. Primers and Gene Identifiers for individual homologues shown can be found in dataset S3. *BnGP4* and *BnEEF* were used as housekeeping genes for qPCR. (C) Seed abortion rates observed in *BnBZIP*_*RNAi*_ siliques. Significance determined using a Student’s T-test (p<0.05). (D-F) Macrographs of siliques of Topas (D) and *BnBZIP11*_RNAi_ (E-F). Aborted ovules are indicated with grey triangles. Scale bar D-E = 5mm. Scale bar F = 1mm.

## Discussion

### The developing *Brassica napus* seed is a resource for examining subgenome expression dominance in neopolyploid development

The phenomenon of progenitor subgenome bias has been observed to varying degree in the vegetative organs of *B. napus* (Chalhoub et al., 2014) and other polyploid plants such as *G. hirsutum* (Flagel et al., 2012; Hovav et al., 2015; Song et al., 2015), *Glycine dolichocarpa* (Coate et al., 2014), *Triticum* (Li et al., 2014), *Arachis* (Clevenger et al., 2016), and *Erythranthe* (Edger et al., 2017). In the older allohexaploid *Triticum aestivum*, and root transcriptomes of *T. aestivum* were dominated by single homoelogues irrespective of subgenome of origin (Leach et al., 2014; Li et al., 2014). Similarly, no overall subgenome bias was reported in spikelet transcriptomes (Wang et al., 2017). Peanut (*Arachis hypogaea* L.) is a relatively recent allopolyploid that also shows minimal expression bias in most tissues; however, there was more prominent expression bias in meristematic or actively growing tissues of *Arachis*, including roots, flowers, shoot tips, and young seeds (Clevenger et al., 2016). Expression bias in *G. hirsutum* seed development was found to be highest during seed elongation and maturation in whole seeds (Hovav et al., 2015), and was found to favour higher expression of its larger *G. arboreum* (A) subgenome over the smaller *G. raimondii* (D) subgenome.

Both tissue and subgenome of origin effects can influence gene expression patterns in the vegetative tissues of *B. napus*, though in both leaves and roots most homoeologous pairs remain unbiased between subgenomes (Chalhoub et al., 2014). Our data show that in whole *B. napus* seeds, subgenome expression bias favours the smaller A^n^ (*B. rapa*) subgenome. The A^n^ subgenome is less densely populated with transposable elements than the C^n^, especially in gene-rich regions. It is possible that epigenetic regulation of transposable elements may affect nearby genes during seed development in *B. napus*. We are currently investigating whole genome methylation patterns and small RNA dynamics in seed development to further address this phenomenon. Taken together, these findings indicate that subgenome expression bias in young allopolyploids is most strongly observed in rapidly growing organs or in tissues undergoing drastic developmental transitions, such as flowers and seeds. The developing *B. napus* seed is therefore a useful resource for interrogating the contributions of subgenomes to a neopolyploid transcriptome during dramatic developmental shifts.

### Asymmetric gene regulation in maternal and filial tissues of the *B. napus* seed

The transcriptional subgenome contributions of an allopolyploid may vary by tissue and developmental stage. A transcriptomic analysis of allohexaploid wheat (*Triticum aestivum* L.) showed significant subgenome expression bias in crown roots (Powell et al., 2017), in contrast to the homoeologue dominance observed when entire root systems are examined (Leach et al., 2014; Li et al., 2014). In allotetraploid *G. hirsutum* seeds, subgenome transcription bias favouring the larger A (*G. arboreum*) is observed in across seed development (Hovav et al., 2015) but if ovule fibres specifically are examined, then higher expression is observed from the smaller D (*G. raimondii*) subgenome (Yoo and Wendel, 2014). The transcriptional profiles of the A and D progenitor subgenomes are distinct in *G*. *hirsutum* ovule fibers (Song et al. 2015), indicating a tissue-specific effect on subgenome expression in the developing seed.

Small subsets of genes also exhibit reversals of expression bias between the shoot and root tissues of *B. napus* (Chalhoub et al., 2014). Our data indicate that bias favouring higher expression of the smaller A^n^ subgenome can be observed in nearly all tissues of the globular stage seed, except the chalazal proliferating tissue. In both Arabidopsis and soybean, it has been suggested that activity of transcription factors plays one of the most important roles in modulating gene expression in developing seeds (Lin et al., 2017). While overall expression bias favoured the A^n^ subgenome in the seed, the A^n^ subgenome transcriptional circuits were enriched primarily in the seed coat and maternal tissues, and the C^n^ subgenome was enriched primarily in the embryo. Interestingly, the chalazal and peripheral endosperm did not contribute to the expression of subgenome-enriched transcription factors in early seed development. This implies that subgenome bias in transcriptional networks is not consistent across the embryo, endosperm, and seed coat of the seed.It is likely that in addition to the action of transcription factors, DNA methylation (Bräutigam and Cronk, 2018) (Bräutigam and Cronk, 2018), translational modifications (Coate et al., 2014), and small RNAs play a role in maintaining differential contributions of subgenomes or specific homoeologues in allopolyploids (Li et al., 2014; Song et al., 2015; Zhang et al., 2018). Given that we observe uneven contribution of transcription factor expression to subgenome-biased coexpression networks, it would be of great value to examine the expression of small RNAs at the whole seed and tissue-specific level in *B. napus*, as has been performed in Arabidopsis (Kirkbride et al., 2019). Such data would provide deeper insight into the contributions of the A^n^ and C^n^ subgenomes to the regulation of gene expression in the allotetraploid *B. napus*.

### Early *B. napus* seed development is regulated by BZIP transcription factors

As proof-of-concept to confirm our predictive regulatory networks, we decided to look at transcriptional networks independent of subgenome of origin in order to get a broader view of the gene regulation in the seed. We examined networks comprising both subgenomes so as to account for any redundancy between homoeologous transcription factors and targets. We performed Fuzzy-K means clustering as a less stringent means of identifying co-expressed gene sets. Early *B. napus* seed development was enriched for Group S1 BZIP transcription factors. nockdown of *BnBZIP11* was sufficient to decrease expression of genes predicted by our network analysis to be targets of genes associated with photosynthesis and cytoskeleton growth leading to a reproductive-lethal phenotype. However, S1 BZIPs tend to act as stronger transcriptional activators as heterodimers (Ehlert et al., 2006), and *BnBZIP11* is co-expressed with two S1 BZIPs, *BnBZIP44*, and *BnBZIP53*, which also show high expression in the chalazal pole and maternal tissues of the globular *B. napus* seed (Ziegler et al., 2019). It would be interesting to see the effect of knockdown of multiple group S1 BZIPs on gene regulation in *B. napus* seed development. Some homologues of the coexpressed BZIPs are expressed outside of the chalazal pole in the endosperm and embryo at the globular stage (Ziegler et al., 2019). A high-resolution spatiotemporal transcriptomic dataset that includes additional developmental stages and that examines the maternal and filial regions of the seed will provide greater insight into how different homoeologues are expressed in different tissues and may lead to novel insight regarding their functions.

## Conclusions

We profiled global shifts in gene activity across the A^n^ and C^n^ subgenomes of allotetraploid seeds of *B. napus*. These data provide valuable insight into the patterns underlying gene expression in both the early and late stages of seed development in one of the most important oilseed crops. Subgenome bias is also observed in predictive regulatory networks of the developing seed transcriptome. The early seed transcriptome is further characterized by uneven contributions from embryo, endosperm, and seed coat to subgenome-biased transcription factor expression. Taken together, gene regulation in the developing *B. napus* seed is a complex landscape defined by subgenome bias and through spatiotemporal regulation of transcription factors.

## Materials and Methods

### Plant growth conditions and sample collection

Transcriptome analyses were performed in *B. napus* DH12075 (whole seed). Plants were grown in growth chambers under long day conditions at 22°C. Flowers were hand-pollinated and collected at 0 days post fertilisation (DPF) (OV), 7DPF (GLOB), 10DPF (HEART), 28DPF (MG), and 35 DPF (DS) between the hours of 14:00-16:00 to minimize time of day effects among samples. Seeds were manually harvested from siliques under RNAase-free conditions and flash-frozen in liquid nitrogen.

### RNA isolation, library preparation, and sequencing

Whole seed tissues were ground in liquid nitrogen. RNA isolation from whole seeds was performed using PureLink™ Plant RNA reagent (Ambion®). RNA quality was assessed using the Agilent 2100 Bioanalyzer. Samples of a minimum RIN of 7 were used for library preparation (Jahn et al., 2008). Library preparation and quality control was performed as in Becker et al. (2017), modified from a high throughput RNA-seq library preparation protocol C2 (Kumar et al., 2012). Three replicates of each stage were prepared, with the exception of the dry seed stage, for which only 2 libraries were prepared as a result of insufficient quantities of RNA for preparation. Libraries were sequenced on the Illumina HiSeq 2500 platform at Genome Quebec (100bp single end reads).

### RNA-seq data analysis

Quality trimming and adaptor removal from sequencing reads was performed using Trimmomatic (Bolger et al., 2014). Read quality was evaluated using FastQC (http://www.bioinformatics.babraham.ac.uk/projects/fastqc/). High quality reads were then aligned to the DH12075 genome (Parkin et al., unpublished) using hisat2 (Kim et al., 2019). Uniquely mapping reads were then passed through to cuffnorm and cuffdiff (Trapnell et al., 2013) to determine transcript accumulation levels and differential gene expression. Transcripts were considered “detected” at FPKM≥5 based on inter-replicate variability was assessed using the CummeRbund suite (Goff et al., 2012). For further differential expression and clustering analyses, a cut-off of FPKM≥10 was selected based onour previous comparisons of RNA-seq and qPCR data (Millar et al., 2015; Ziegler et al., 2019).Global clustering of transcripts (FPKM≥10) was performed using pvclust (https://CRAN.R-project.org/package=pvclust).

Based on this cutoff (FPKM≥10), transcripts were screened for accumulation intersection between seed stages (OV, GLOB, HEART, MG, DS) and phases (early = OV, GLOB, HEART; late/maturation = MG, DS; morphogenesis = GLOB, HEART) and visualized using the UpSetR package (Conway et al., 2017). We also performed Fuzzy-K means clustering as a less stringent means of identifying co-expressed genes. Temporal patterns of transcript accumulation were found using a Fuzzy K means clustering algorithm (http://cran.r-project.org/web/packages/cluster. Homologous genes in the Darmor Bzh annotation were identified, and enrichment of transcript accumulation patterns was performed using SeqEnrich (Becker et al. 2017).

### Accession Numbers

All data have been deposited at the Gene Expression Omnibus (GEO): GSE144771.

### Generating *BnBZIP11* knockouts

Knockdown of transcription factors was performed in *Brassica napus* cv. Topas using RNA interference. A 489bp fragment of high homology between all expressed *BnBZIP11* homologous transcripts was identified and cloned from seed cDNA. The *BnBZIP11* fragment was cloned into the Gateway cassette of pENTR4 (Invitrogen®). The fragment was sequenced to confirm its identity (Dataset S3). Gateway cloning was used to transfer the fragment into the two intron hairpin GW sites of pHELLSGATE8 (Helliwell et al. 2003). Successful transfer into destination vectors was confirmed by PCR and restriction digest analysis. Expression constructs were transferred into *Agrobacterium* GV3101. Generation of transgenic *B. napus* was performed using cotyledon transformations according to Bhalla and Singh (2008). Successful transformation events were selected with kanamycin and confirmed using PCR. Transcript knockdown was confirmed in T0 and T1 plants using qPCR (qScript cDNA synthesis kit, QuantaBio®, and Luna qPCR mix; NEB®) according to manufactuers’ protocols. The efficacy of *BnBZIP11* knockdown was variable in subsequent generations of transgenic plants, so analysis was performed in T1 plants. Since relatively high expression of an intron hairpin construct is required for effective transcript knockdown, and high expression of a transgene can lead to its silencing (Schubert et al., 2004), we recommend a different approach such as inducible transgene expression in the future.

### Gene network validation

Primers were designed against target genes from the GO terms of the regulatory network, and knockdown of these target genes in the *BnBZIP11* knockdown mutant were confirmed using qPCR in siliques (10DPF, corresponding to the heart stage in morphogenesis). *BnGP4* and *BnEEF* were used as housekeeping genes for comparison. Primers and full gene IDs of all transcripts examined can be found in Dataset S3.

## Acknowledgments

We thank Dr. Steve Whyard (University of Manitoba) for the pHELLSGATE8 construct, and Dr. Dana Schroeder for the *Agrobacterium* GV3101 cells. We thank Drs. Genyi Li and Claudio Stasolla, Kenny So, and Hanna Dandena (University of Manitoba) for their input on *Brassica* transformation and tissue culture.

## Supplemental Figure Legends

**Figure S1.** Number of transcripts expressed at a low (purple, 5<FPKM<10), medium (teal, 10<FPKM<25) and high (green, FPKM>25) levels throughout seed development.

**Figure S2.** Heatmap some of the most highly expressed genes in the dataset. FPKM values have been log_2_ transformed.

**Figure S3.** Predictive regulatory network of *B. napus* seed development at the MG stage. The network shows transcription factors (purple rectangles), DNA binding motifs (teal hexagons) to genes that below to a biological processes (blue circles) within a co-expressed gene set (gray diamond). (A) Network of MG stage coexpressed genes originating from the A^n^ subgenome. (B) Network of MG stage coexpressed genes originating from the C^n^ subgenome. Gene lists and enrichment outputs can be found in Dataset S2.

## Supplemental Datasets

Dataset S1. Transcript levels of all detected transcripts in the developing seed (OV, GLOB, HEART, MG, DS), including lists of differentially expressed and stage and phase-specific genes. Differentially expressed genes were identified using Cuffdiff. Stage- and phase-specific transcripts (FPKM>10 in stage/phase, FPKM<10 in other stages/phases) are listed, with subgenome of origin indicated.

Dataset S2. GO and network enrichment outputs for the stage and phase specific gene sets highlighted in Figures 4 and S8 are included. We also performed Fuzzy K means clustering across the A^n^ and C^n^ subgenomes inclusively. Gene lists and pattern outputs are indicated, along with GO enrichment and network outputs for all analyses.

Dataset S3. Validation of *BnBZIP11* knockdown. The sequence cloned into the RNAi hairpin vector as well as the primers used to validate the expression of *BnBZIP11* and its targets are included.

